# SMER28 attenuates PI3K/mTOR signaling by direct inhibition of PI3K p110 delta

**DOI:** 10.1101/2021.12.10.471916

**Authors:** Marco Kirchenwitz, Stephanie Stahnke, Silvia Prettin, Malgorzata Borowiak, Carmen Birchmeier, Klemens Rottner, Theresia E.B. Stradal, Anika Steffen

**Author notes:** Authors for correspondence: Anika Steffen or Theresia E.B. Stradal, Anika Steffen, Theresia E.B. Stradal.

## Abstract

SMER28 (Small molecule enhancer of Rapamycin 28) is an autophagy-inducing compound functioning by a hitherto unknown mechanism. Here we confirm its autophagy-inducing effect by assessing classical autophagy-related parameters. Interestingly, we also discovered several additional effects of SMER28, including growth retardation and reduced G1 to S phase progression. Most strikingly, SMER28 treatment led to a complete arrest of receptor tyrosine kinase signaling, and consequently growth factor-induced cell scattering and dorsal ruffle formation. This coincided with a dramatic reduction of phosphorylation patterns of PI3K downstream effectors. Consistently, SMER28 directly inhibited PI3Kδ and to a lesser extent p110γ. The biological relevance of our observations was underscored by interference of SMER28 with InlB-mediated host cell entry of *Listeria monocytogenes*, which requires signaling through the prominent receptor typrosine kinase c-Met. This effect was signaling-specific, since entry of unrelated, gram-negative *Salmonella* Typhimurium was not inhibited.

## Introduction

Autophagy is an evolutionarily highly conserved cellular recycling process that is modulated by signals such as nutrient deprivation, metabolic stress, energy depletion and hypoxia. All these signals regulate induction of autophagy, which also occurs on a basal level as a homeostatic process. A specific machinery controls this catabolic process in which double-membrane vesicles are formed to enclose various cytoplasmic constituents such as protein aggregates, damaged organelles or intracellular pathogens. These double-membraned structures are termed autophagosomes and are finally delivered to and fuse with lysosomes to degrade and recycle the basic constituents (1-3). A cellular kinase scaffold termed mTORC1 complex is central to the regulation of autophagy and integrates signals on oxidative and energy stress, but also on the availability of e.g. growth factors or insulin to balance cellular growth. mTORC1 shares the protein subunits mTOR, Deptor and mLST8 with the mTORC2 complex, but is distinct from mTORC2 through harboring the subunit RAPTOR. In addition, mTORC1 activity is selectively inhibited by rapamycin through binding of a FKBP12-rapamycin complex. In contrast, mTORC2 retains its kinase activity upon acute rapamycin treatment, likely because the mTORC2-specific subunit RICTOR sterically prevents FKBP12-rapamycin binding (4). A selective activator of mTORC1 activity is the small GTPase Rheb, which in turn is regulated by PI3K, AKT and the Rheb1-deactivating TSC complex. Thus, growth factor signaling through PI3K leads to active, GTP-bound Rheb1, hence activating mTORC1 (1). However, the enzymatic activity of PI3K is not only central to growth factor signaling, but may also cause malignant transformation of various cell types if the activity overshoots (5). The etiology of hyperactivation can be due to copy number gain, increased expression or mutations leading to pronounced or constitutive activation of PI3K isoforms, as well as loss of antagonist function such as of the lipid phosphatase PTEN (6). Interestingly, a growing body of evidence shows that the PI3K/AKT pathway is interconnected with autophagy, suggesting that this might be a sensitive axis to interfere with cancer progression. In fact, a dual role of autophagy in the initiation and development of cancer currently constitutes an active area of research. On one hand, autophagy is induced in different tumor types as a response mechanism to therapy. In line with this observation, inhibition of autophagy was found to render cancer cells vulnerable for therapy (7). On the other hand, downregulation of autophagy by different means was shown to augment tumor development and, *vica versa*, induction of autophagy conditions supports defeating of tumors. More specifically, deletion of several ATG (autophagy related genes) and pro-autophagic kinases such as AMBRA1 and Beclin was observed to lead to the appearance of tumor lesions (8-12). In addition, drugs known to target the PI3K/AKT/mTOR axis were reported to enhance radiation sensitivity in cancer cell lines and cancer models in pre-clinical studies (13-15). Indeed, screening of new target compounds and drug combinations already revealed anticancer effects by attenuating this PI3K/AKT/mTORC1 signaling axis (16-18).

SMER28 was identified in a screen of autophagy enhancing compounds acting independently of rapamycin (19). While mTORC1 is clearly central to the regulation of autophagy, and rapamycin suppresses the kinase activity of mTORC1 through allosteric inhibition, the mode of action of SMER28 has remained elusive. Here we reveal that SMER28 induces autophagy through a hitherto unknown mode of action. We show that SMER28 directly inhibits PI3K by binding to its catalytically active subunit p110. Moreover, we provide evidence that SMER28 causes growth retardation, accompanied by a partial arrest of the cell cycle in G1. SMER28 has little effect, however, on cell viability as evidenced by cell cytotoxicity measurements. Attenuation of PI3K signaling by SMER28 is further confirmed by strongly reduced phosphorylation levels of Thr308-Akt and Ser473-Akt, two target sites of active PI3K. Consequently, SMER28 effectively blocked growth factor signaling downstream of receptor tyrosine kinases, as exemplified by the complete abolishment of cell scattering and dorsal ruffle formation elicited by HGF or PDGF. Together, we here unveil the mechanism of autophagy induction by SMER28 through its direct targeting of the PI3K/AKT signaling axis, suggesting it as promising lead structure for the development of anti-cancer drugs, most specifically B cell lymphomas due to its high affinity for the p110δ subunit.

## Results

### SMER28 moderately increases autophagy

In order to document the effects of SMER28 on the regulation of autophagy marker proteins, we set out to apply a series of analyses. We first assessed the appearance of autophagosomes in control and SMER28-treated U-2 OS cells by immunofluorescence with antibodies directed against LC3 and SQSTM1/p62 (Sequestosome-1; hereafter p62), two classical markers of autophagosomes (20). Antibody stainings revealed a higher number of LC3 and p62-positive autophagosomal puncta upon treatment with SMER28 (50 µM, Figure 1A). Quantification showed a comparable increase in numbers of LC3- and p62-positive puncta after SMER28 treatment (Figure 1B). We also measured the total area of LC3 and p62 puncta per cell and per autophagosome. We found that SMER28 treatment increased the area of LC3 and p62 per cell approximately two to threefold (Figure 1C), while the average size of each autophagosome was only slightly augmented (Figure 1-figure supplement 1A), indicating an increase in the number of autophagosomes rather than an enlargement of these structures once formed. In addition, we found elevated levels of LC3-II, the lipidated form of LC3 (21), in SMER28-treated samples by Western blotting (Figure 1D), confirming earlier observations (19). Surprisingly however, levels of p62 were also moderately increased after treatment with 50 µM SMER28 (Figure 1E). The induction of autophagy is usually connected to p62 degradation, since it links polyubiquitinated proteins to the autophagosome and hence eventually to autophagic digestion. Thus, we investigated whether SMER28 might also have an inhibitory effect on the proteasomal pathway, which would explain the observed elevated p62 levels. To compare the effect of SMER28 treatment to well-established proteasomal inhibitors, we included bortezomib and MG-132 (22) in our analyses. While treatment of cells with bortezomib and MG-132, respectively, led to an approximately 1.5-fold increase in p62 levels, SMER28 had a comparably mild effect (Figure 1E). Similarly, treatment with bortezomib and MG-132, respectively, increased LC3-II levels to a much higher extent than treatment with SMER28 (Figure 1D). The induction of autophagy by proteasomal inhibitors such as MG-132 has been described earlier (23, 24). Combining SMER28 with these proteasomal inhibitors resulted in lower LC3-II and p62 levels as compared to proteasome inhibition by MG-132 and bortezomib, respectively, alone (Figure 1D, E). This suggests that SMER28 does not inhibit the proteasome, but instead may stimulate the turn-over of autophagy-related marker molecules. Interestingly, another, well-established autophagy-inducing drug, rapamycin (25) showed similar effects. Combined treatment of rapamycin together with MG-132 and bortezomib, respectively, reduced the average levels of LC3-II to extents similar to SMER28 (Figure 1D). To exclude that the observed increase of LC3-II and p62 is simply due to impaired lysosomal function, we compared the levels of LC3-II and p62 after SMER28 treatment alone with combined application with bafilomycin A1, a specific v-ATPase inhibitor that inhibits the acidification of normally acidic organelles such as lysosomes (26). The endogenous LC3-II and p62 levels were substantially increased after co-treatment with SMER28 and bafilomycin A1 as compared to SMER28 treatment alone (Figure 1D, E), suggesting that SMER28 does not simply block lysosomal degradation. In an independent assay, we tested whether application of SMER28 leads to an accumulation of polyubiquitinated proteins. While MG-132 and bortezomib clearly increased the levels of high molecular weight polyubiquitinated proteins, control and SMER28-treated cells showed indistinguishable levels of polyubiquitinated proteins (Figure 1 – Figure supplement 1B).

**Figure 1.**
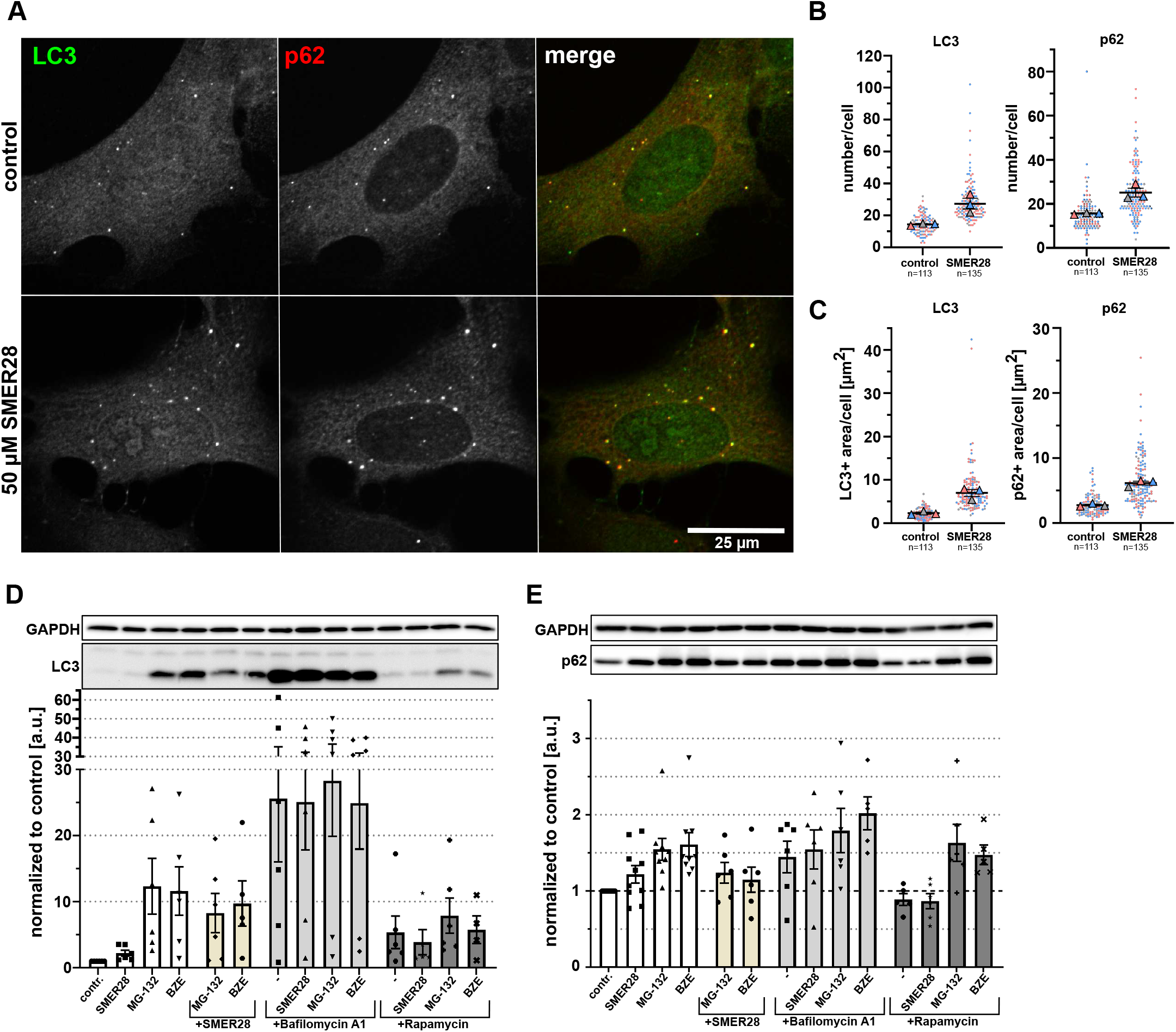
SMER28 stimulates autophagy in U-2 OS cells. **(A)** U-2 OS cells were left untreated (upper panel) or treated with 50 µM SMER28 for 16 h (lower panel). Cells were stained with p62/SQSTM1 and LC3 antibodies to assess markers for autophagosomes and visualized by confocal spinning disk microscopy. Merged images show LC3 in green and p62 in red. **(B-C)** Quantification of average numbers (B) and total areas (C) of LC3-positive and p62-positive puncta per cell. Data are shown as scatter plots with means ± s.e.m.; n= total number of cells analyzed. **(D-E)** U-2 OS cells were treated with 50 µM SMER28, 1 µM MG-132, 15 nM bortezomib (BZE) and in the presence or absence of 250 nM bafilomycin A1 or 300 nM rapamycin for 16 h, as indicated. Protein levels of LC3 (D) and p62 (E) were assessed by WB; GAPDH was used as loading control. Graphs show quantifications of relative LC3-II and p62 levels after treatments as indicated. Data are means ± s.e.m; n≥5.

### SMER28 treatment leads to growth arrest

During our experiments, we noticed retarded cell growth in SMER28-treated conditions. To quantify this effect, we measured cell confluence of SMER28- and rapamycin-treated cells compared to controls over a time period of 47 hours. We found that 50 µM SMER28 retarded cell growth to extents comparable to 300 nM rapamycin (Figure 2A-B). We also tested increasing concentrations of SMER28 (not shown) and found that 200 µM led to almost complete growth arrest after an initial lag phase of approximately 8 hours, during which cell growth could still be observed (Figure 2A). In order to compare this cytostatic effect to well-established cytostatic drugs, we compared the effect of SMER28 and rapamycin to paclitaxel and epothilone B (27, 28) on cell confluence and cell death using PI (propidium iodide)-staining. Interestingly, the cytotoxic effects of 50 µM SMER28 and 300 nM rapamycin were similarly low (both below 1% PI-positive area per total cell area), whereas the cytostatic effect of 200 µM SMER28 was almost as strong as application of paclitaxel and epothilone B, respectively (Figure 2B). This effect was accompanied by an increased portion of dead cells, expressed as PI-positive area per total cell area (Figure 2B), indicating that the stronger growth inhibition upon increased SMER28 concentrations correlates, at least in part, with accelerated cell death. This is comparable to the cytotoxic effects of paclitaxel and epothilone B, which are known to arrest the cell cycle at the G2/M transition through stabilization of microtubules (29). We thus wondered whether SMER28 could have similar effects and analyzed the cell cycle with different concentrations of SMER28- and epothilone B-treated cells by propidium iodide staining. Surprisingly, we found that the percentage of cells in G1 expanded with increasing SMER28 concentrations (Figure 3A, B), suggesting that SMER28 arrests cells at least in part at the restriction point. The cell cycle is tightly regulated by the activity of cyclin dependent kinases (Cdks), which in turn are regulated by the abundance of their inhibitors and the cyclins. Thus, we analyzed whether SMER28 affects the protein levels of aforementioned factors. CDK4/6 heterodimerize with D cyclins to form active kinase complexes allowing cell cycle progression from G1 to S, while the activity of these kinase complexes is inhibited by association of p27/Kip and p21 Waf1/Cip1 with CDKs and cyclins through formation of heterotrimeric complexes (30). CDK4 and CDK6 levels were found to be largely unchanged in samples of control as well as SMER28-treated cells after 24 h (Figure 3C). Interestingly however, both cyclins D1 and D3 showed decreased protein levels after 24 h incubation with 50 µM and 200 µM SMER28 (Figure 3D). Furthermore, expression of the inhibitory subunit p27 was not affected by SMER28 treatment, while p21 levels decreased under SMER28 conditions (Figure 3E). This suggests that the overabundance of p27 during the concomitant lack of the activating subunits cyclin D1 and D3 may lead to the observed partial arrest in G1 to S phase.

**Figure 2.**
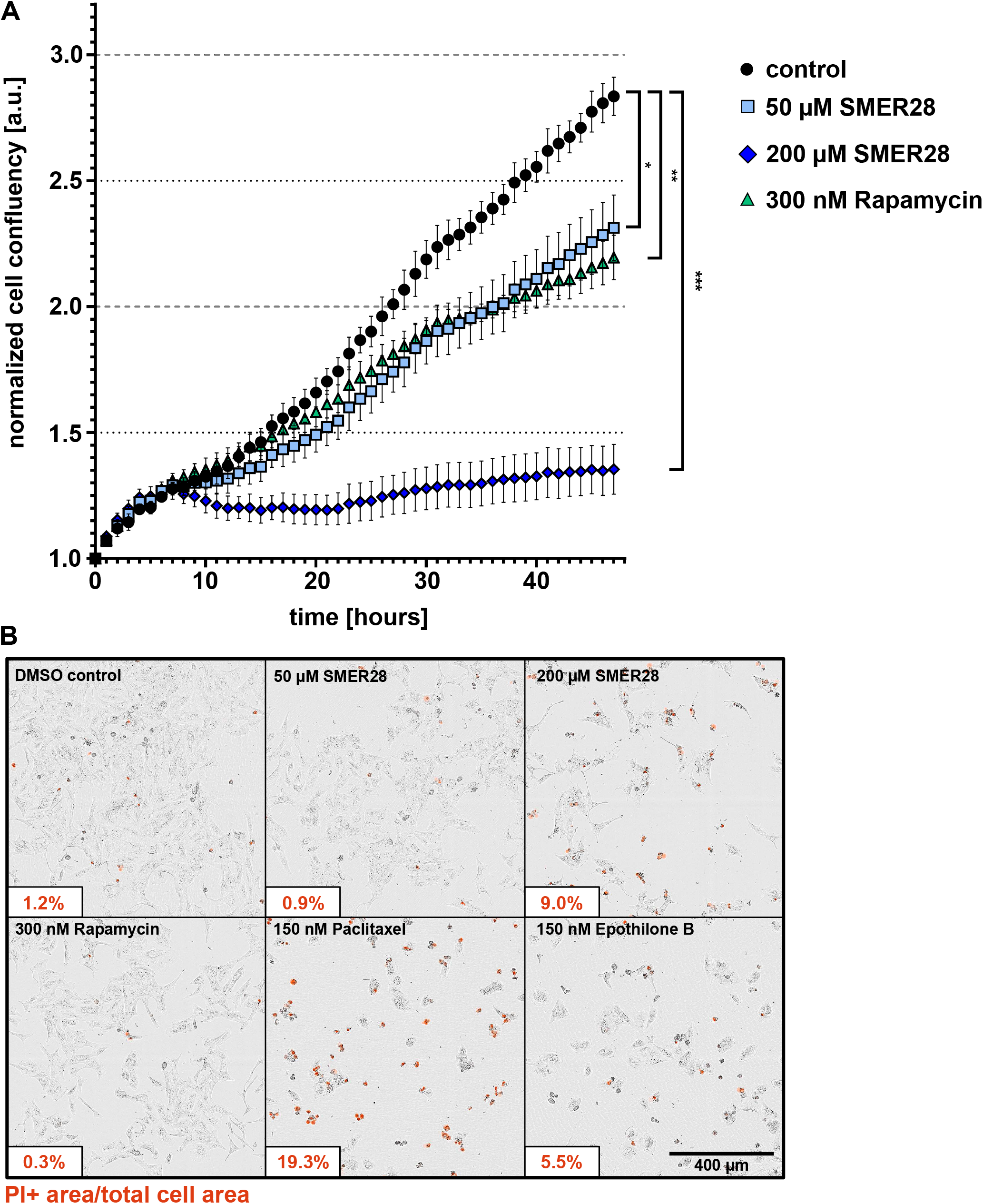
SMER28 treatment inhibits cell proliferation in a dose-dependent manner. **(A)** U-2 OS cells were treated as indicated and immediately recorded by phase contrast imaging for 47 hours. The graph shows the mean confluence ± s.e.m. from at least four independent experiments with ≥4 replicates. *p<0.05, **p<0.01, ***p<0.001, two-way ANOVA. **(B)** U-2 OS cells were treated as indicated, stained with propidium iodide (PI) and imaged by phase contrast and fluorescence microscopy. Images show overlays after 47 h. Insets each display PI (propidium iodide)-positive area per total cell area in percent. Data are means from three independent experiments with at least four replicates each.

**Figure 3.**
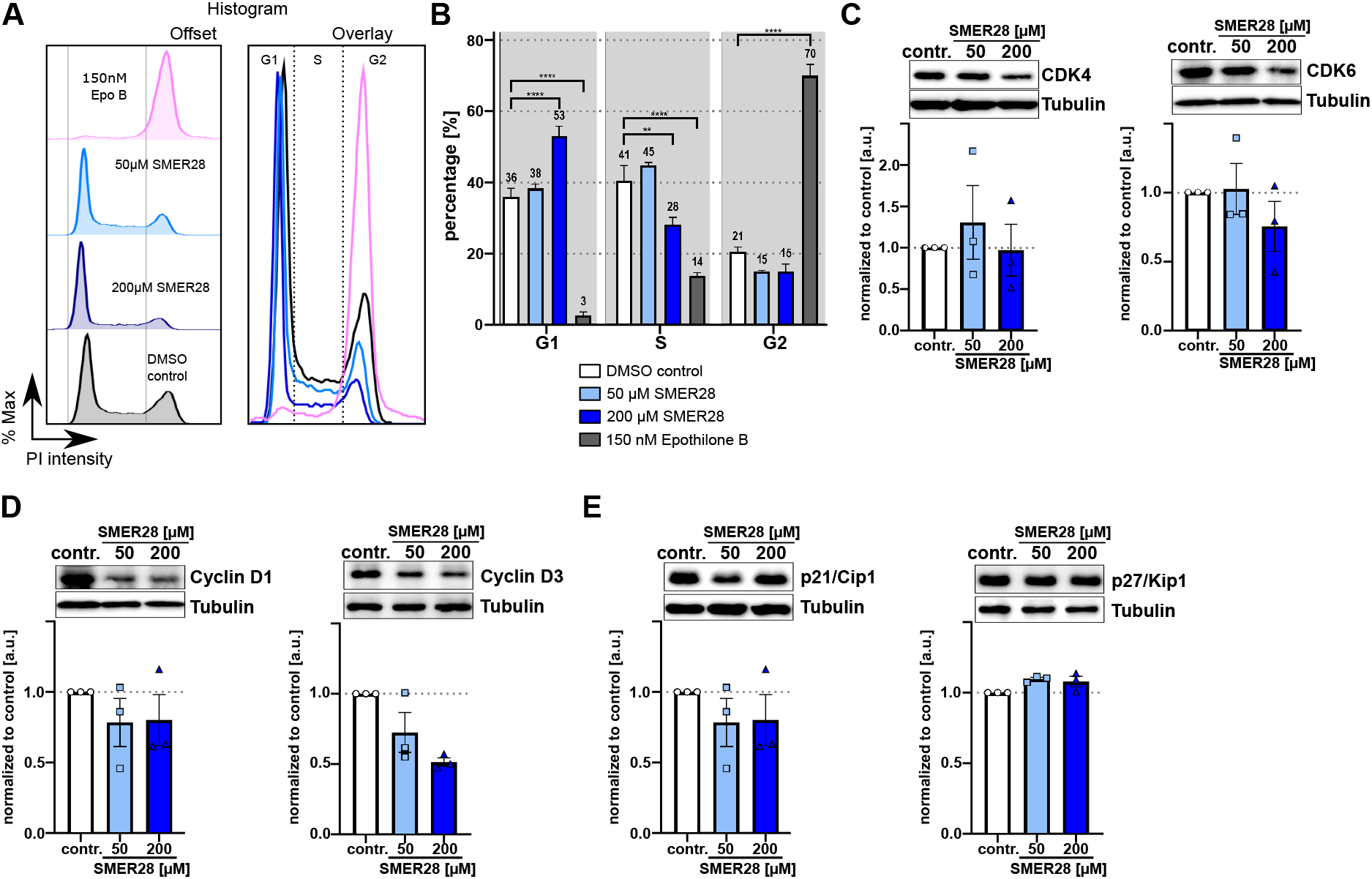
SMER28 treatment induces G1 cell cycle arrest. **(A-B)** Flow cytometry analysis of the cell cycle distribution using PI (propidium iodide) staining after 24 h incubation of U-2 OS cells with different concentrations of SMER28 or epothilone B, as indicated. **(A)** Representative histograms depicting the distribution of cell cycle phases for indicated treatments. Histograms are shown as offset (left) and overlay (right). **(B)** Bar plots showing the respective percentage of cell cycle phases (G1, S and G2) according to PI intensity. Percentage of each cell cycle phase was calculated by Watson Pragmatic model using FlowJo software. Data are means from at least three independent experiments ± s.e.m. *p<0.05, ****p<0.0001, two-way ANOVA. **(C-E)** U-2 OS cells treated with 50 or 200 µM of SMER28 were harvested after 24h. Protein expression levels of CDK4 and CDK6 **(C)**, Cyclin D1 and Cyclin D3 **(D)** as well as p21/Cip1 and p27/Kip1 **(E)** were assessed by WB. Bar graphs show protein levels calculated relative to tubulin. Data are means ± s.e.m; n=3.

### SMER28 interferes with receptor tyrosine kinase signaling

Next, we addressed which upstream regulatory pathway is targeted by SMER28 that may eventually lead to growth arrest. Signaling of growth factor receptor tyrosine kinases (RTKs) is well established to be connected to proliferation as well as autophagy (31). We thus tested whether SMER28 affected epithelial cell scattering induced by HGF (hepatocyte growth factor) by recording MDCK cells over a time period of 12 hours of stimulation with 20 ng/ml HGF. DMSO control-treated MDCK cells scattered markedly from pre-formed colonies (Figure 4A, left panel, 4B; see also Supplementary movie 1). In stark contrast, application of 50 µM and 200 µM SMER28, respectively, completely abrogated scattering of MDCK colonies (Figure 4A, middle and right panel, 4B; see also Supplementary movie 1), indicating a strong defect in signal transduction downstream of the HGF/c-Met receptor. In a complementary approach, we recorded growth factor responsiveness by stimulating serum-starved NIH3T3 cells with either HGF or PDGF (platelet derived growth factor). F-actin stainings revealed prominent responses in control cells upon treatment with HGF or PDGF (Figure 4C, upper panel (32, 33)). Indeed, more than 50% of control cells responded with well-developed, dorsal ruffles upon treatment with either growth factor (Figure 4C, D). In contrast, less than 12% (50 µM) and 6% (200 µM) of SMER28-treated cells displayed dorsal ruffles upon HGF or PDGF stimulation, whereas the portion of cells with peripheral ruffles was nearly the same in all conditions (Figure 4C, D). Of note, most of the dorsal ruffles still formed in SMER28-treated cells were much smaller and/or not completely circularly closed. Collectively, these data reveal that SMER28 severely impairs signaling downstream of RTKs such as HGF and PDFG.

**Figure 4.**
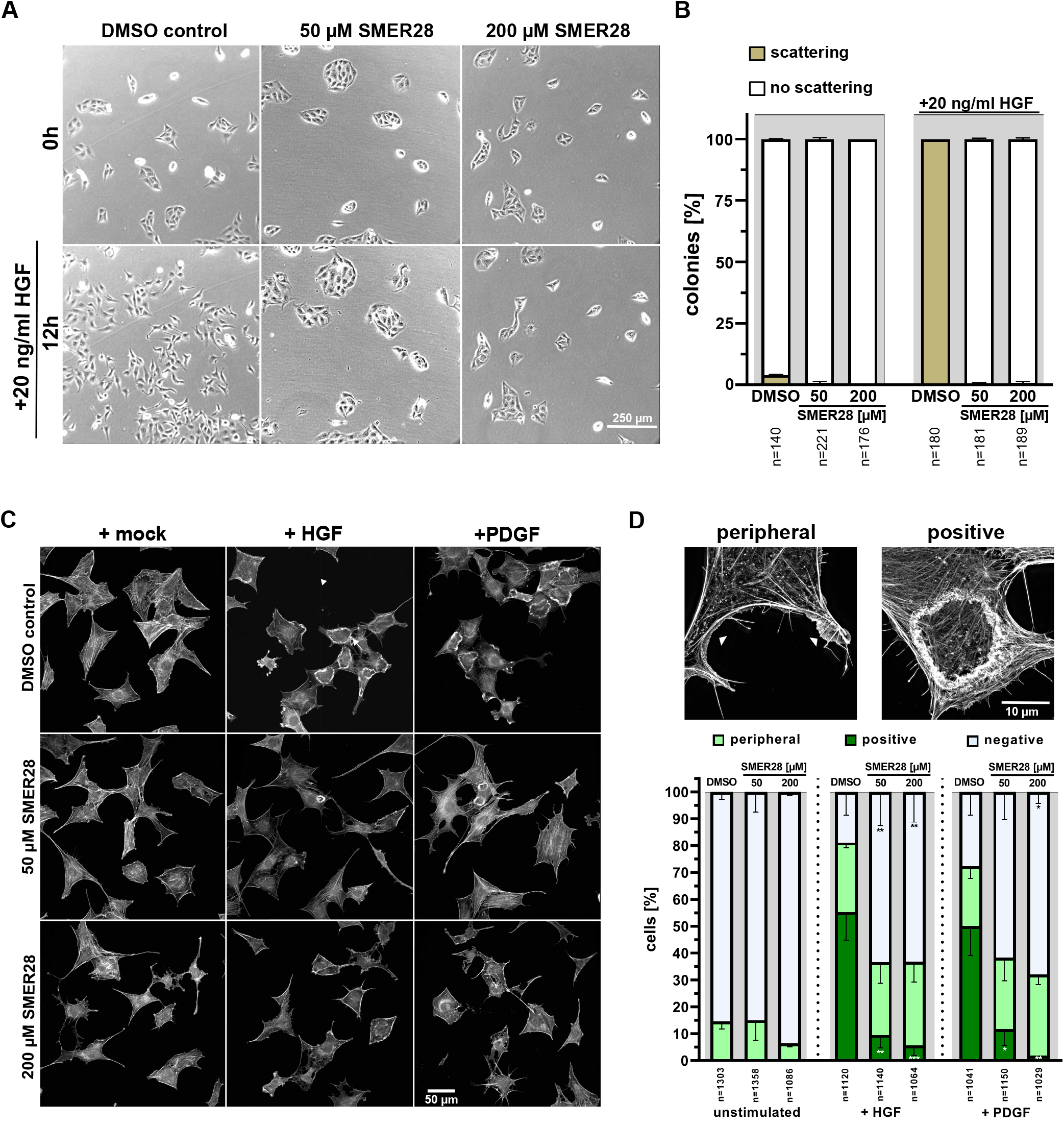
SMER28 inhibits growth factor-mediated cell scattering and membrane ruffling. **(A)** Colony-forming MDCK cells were stimulated with 20 ng/ml HGF in the presence or absence of 50 or 200 µM SMER28, as indicated, and imaged every 15 minutes by phase contrast microscopy for 12h. **(B)** Quantification of MDCK scatter assay. Colonies were categorized as “no scattering” versus “scattering” as described in Materials and Methods. Data were collected from three independent experiments and represent means ± s.e.m; n=total number of cells analyzed. **(C)** NIH/3T3 cells were serum-starved for 16h in DMEM with treatments as indicated on the left. Cells were then treated with DMEM with or without HGF or PDGF for 5 min, as indicated, stained with phalloidin to visualize the F-actin cytoskeleton and imaged by spinning disk microscopy. **(D)** Quantification of growth factor responses according to examples shown above the graph. Images show maximum intensity projections of super-resolution SIM images. Cells were categorized as being non-responsive (negative) *versus* harboring peripheral ruffles (peripheral) or circular dorsal ruffles (positive). Data were collected from three independent experiments and represent means ± s.e.m; n=total number of cells analyzed. *p*<*0.05, **p*<*0.01, ***p<0.001, one way ANOVA.

### SMER28 directly targets PI3K subunits gamma and delta

All findings described so far pointed towards a direct effect of SMER28 on the RTK/PI3K/mTOR signaling axis, since we and others had observed an impact on autophagic flux (Figure 1) (19, 34), and, in addition, on RTK signaling and growth retardation (see Figures 2-4). The progression of cancer is frequently accompanied by aberrant kinase activity in this particular signaling pathway. To determine the molecular impact on this signaling cascade in more detail, we compared the effects of 50 µM and 200 µM SMER28 with rapamycin, a direct inhibitor of mTOR. Rapamycin affected phosphorylation of mTOR at Ser2448 as well as its downstream substrate p70S6K (p-Thr389) (Figure 5A, B). 50 µM SMER28 did not change phosphorylation levels of mTOR (p-Ser2448) after 4 hours of treatment, confirming initial observations on rapamycin-independent effects of SMER28 (19). Surprisingly however, 200 µM SMER28 reduced the levels of phosphorylated mTOR to extents comparable to rapamycin (Figure 5A, B). This SMER28-dependent reduction of mTOR phosphorylation is also recapitulated by phosphorylation levels of p70S6K (p-Thr389) being reduced by approximately half (Figure 5A, B). While mTORC1 regulates various metabolic processes in cells, ULK1 is the first central downstream kinase specific for autophagy. One critical residue in ULK1 is Ser758 (Ser757 in mouse), which is phosphorylated by active mTORC1 to deactivate ULK1 (35). After 4 hours of treatment with 50 µM SMER28, we found a slight reduction of Ser758 phosphorylation of ULK1 (Figure 5A, B). Of note, 200 µM SMER28 treatment reduced the levels of p-Ser758 ULK1 by half, while rapamycin showed a less severe effect (28% reduction). This rather mild effect of rapamycin on the Ser758 phosphorylation site of ULK1 is known to be due to the low sensitivity of the mTORC1 phosphorylation site to rapamycin (36). Together, the effects of SMER28 on these signaling kinases may account for the enhanced autophagy observed above (Fig. 1) (19).

**Figure 5.**
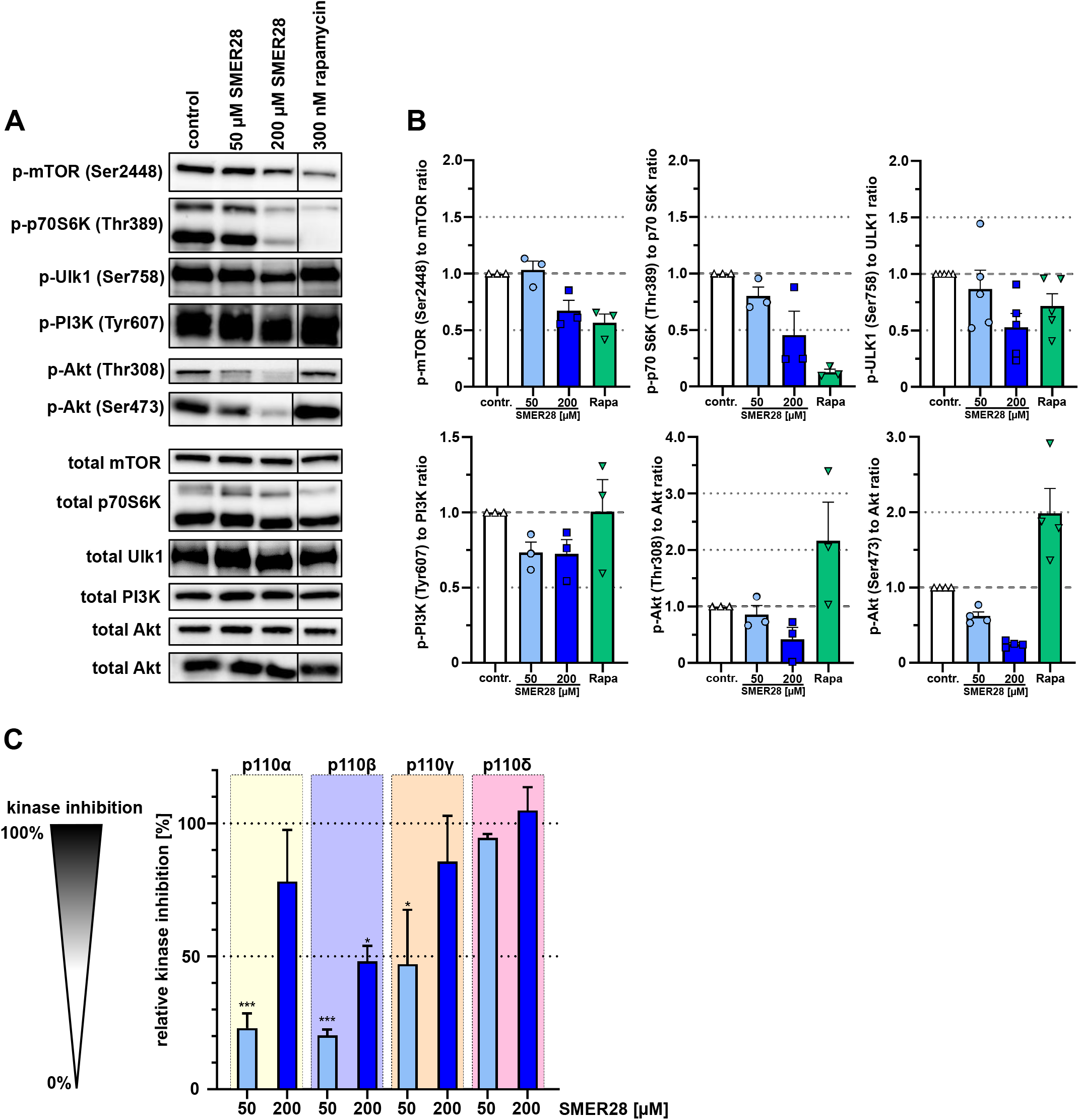
SMER28 inhibits PI3K signaling. **(A)** Western Blot analysis of U-2 OS cells in the presence or absence of 50 or 200 µM of SMER28 or 300 nM Rapamycin (Rapa), as indicated, for 4h were analyzed for levels of phospho-proteins and their respective total proteins. **(B)** Quantifications of relative ratios of phosphorylated proteins to total proteins as indicated. Data are means ± s.e.m. of protein levels derived from at least three independent experiments. **(C)** Inhibition of activities of class I PI3 kinases containing the regulatory subunits p110α, -β, -γ and -δ by 50 or 200 µM SMER28, as measured using an *in vitro* PI3 kinase activity/inhibitor ELISA. The respective relative kinase inhibition was normalized to the data of the reaction without kinase, equaling 100% inhibition per definition. Data represent mean values ± s.e.m; n=4. *p<0.05, **p<0.01, ***p<0.001, compared to the reaction without kinase (not shown, 100%); one way ANOVA.

Since we had observed growth retardation through G1 cell cycle arrest and defective RTK signal transmission, we addressed whether SMER28 specifically affects kinase activity upstream of mTOR. Surprisingly, we found that SMER28 mildly reduced the levels of pTyr607 PI3K, whereas rapamycin did not change these levels, as expected (Figure 5A, B). The oncogene AKT is an indirect downstream effector of PI3K and phosphorylated by mTORC2 at Ser473 (37, 38) and by PDK1 at Thr308 (39). We thus quantitatively analyzed both phosphorylation sites of AKT and found, to our surprise, strongly reduced levels of Ser473-AKT phosphorylation by SMER28 (38% and 75% reduction with 50 µM and 200 µM SMER28, respectively) (Figure 5A, B). In contrast, rapamycin increased p-Ser473-AKT almost twofold (Figure 5A, B), which has been described to occur by mTORC1 inhibition (40, 41). These data thus corroborate the mTORC1-independent mode of action of SMER28 (19). Interestingly, the levels of phosphorylated AKT at residue Thr308 were reduced by more than 50% by 200 µM SMER28, and to much lesser extent with 50 µM SMER28 (Figure 5A, B), indicating that it acts on AKT phosphorylation indirectly and thus independent of its specific kinases mTORC2 and PDK1. We tested therefore whether SMER28 compromises PI3K function directly using an *in vitro* kinase assay. The active enzyme PI3K converts PIP2 to PIP3, which can be quantitatively assessed by adding biotinylated PIP3 (B-PIP3) as tracer. Class I PI3Ks comprise four different members that are differentiated by their catalytic subunits p110 alpha, beta, gamma and delta (5). The ubiquitously expressed subunits alpha and beta were markedly inhibited with 200 µM SMER28, but only mildly affected with 50 µM of the compound. Notably however, PI3K activity exerted by the p110 delta subunit was eliminated with 200 µM SMER28 and almost entirely suppressed (by 87%) with 50 µM SMER28 (Figure 5C). The activity of the gamma subunit was inhibited more moderately, i.e. by 43% and 86% with 50 µM and 200 µM SMER28, respectively (Figure 5C). To confirm the mode of action of SMER28 in a cellular assay, and to assess the relevance of this drug in *in vivo* applications, we investigated the effect of SMER28 on *Listeria* invasion. *Listeria monocytogenes* is a gram-positive, bacterial pathogen that can cause abortions and illnesses such as meningitis. *Listeria* invades host cells by binding of its surface and/or secreted factors, internalin A (InlA) and InlB to the host cell receptors cadherin-1/E-cadherin and c-Met/HGFR, respectively. As the InlA-E-cadherin interaction is restricted to the human system (42), internalin-specific entry into murine cells depends entirely on the InlB-Met pathway. Met/HGFR is widely expressed; so signaling can be induced by binding of its natural ligand HGF in various cell lines (see also Figure 4) as well as by binding of bacterial InlB, known to involve PI3K activation (43-46). We therefore addressed the invasion capability of *Listeria* in the presence and absence of SMER28 and compared this with the non-specific invasion assessed by *Listeria* lacking both InlA and –B (ΔInlA/B). Strikingly, 50 µM SMER28 reduced *Listeria* wildtype invasion by almost 90%, while 200 µM SMER28 completely abolished entry. Wortmannin and LY294002, two different pan-PI3K inhibitors, reduced *Listeria* invasion by only 39% and approx. 75%, respectively (Figure 6A, B). The differences of effects of both inhibitors could be due to the poor stability of wortmannin in solution (47). Additional effects seen with these inhibitors on the invasion of Internalin A/B deletion mutants (*Listeria* ΔInlA/B) indicated a role for PI3K on non-specific, i.e. InlB-independent entry (Figure 6A, C). This conclusion was corroborated by comparing *Listeria* wildtype entry (with or without PI3K inhibition) into cells with or without genetic elimination of the HGF/InlB-receptor (c-Met). Results from c-Met-KO cells (Figure 6C) essentially phenocopied *Listeria* ΔInlA/B data in wildtype fibroblasts (Figure 6A, B). Moreover, wortmannin was again less effective in control and c-Met KO cells than SMER28 at both concentrations employed (Figure 6C). These data demonstrated that SMER28 has robust effects on InlB-mediated *Listeria* invasion known to involve PI3K function in different cells lines. By contrast, gram-negative *Salmonella enterica* Serovar Typhimurium causing food-borne diarrhea invade host cells largely independently of class I PI3 kinases (48, 49). We thus also explored the invasion capacity of *Salmonella* in the presence and absence of SMER28. Strikingly, SMER28 treatments did not significantly affect the invasion capacity of *Salmonella* into NIH/3T3 fibroblasts (Figure 6D), similar to what was found with LY294002 (Figure 6E). To exclude that the compound has a harmful effect on *Listeria* itself, we performed growth curves of *Listeria* in bacterial growth and cell culture medium supplemented with 50 µM and 200 µM SMER28, respectively, or gentamycin as a classical antibiotic as control, and compared these growth curves to DMSO control. As shown in Figure 6F, none of the SMER28 conditions led to an adverse growth affect, while gentamycin effectively inhibited *Listeria* growth, excluding that the reduced *Listeria* invasion rates observed above are due to toxic effects of SMER28 on the bacteria. Together, our data reveal a specific inhibition of class I PI3 kinases by SMER28 with differential activity towards respective p110 subunits, in particular p110 delta.

**Figure 6.**
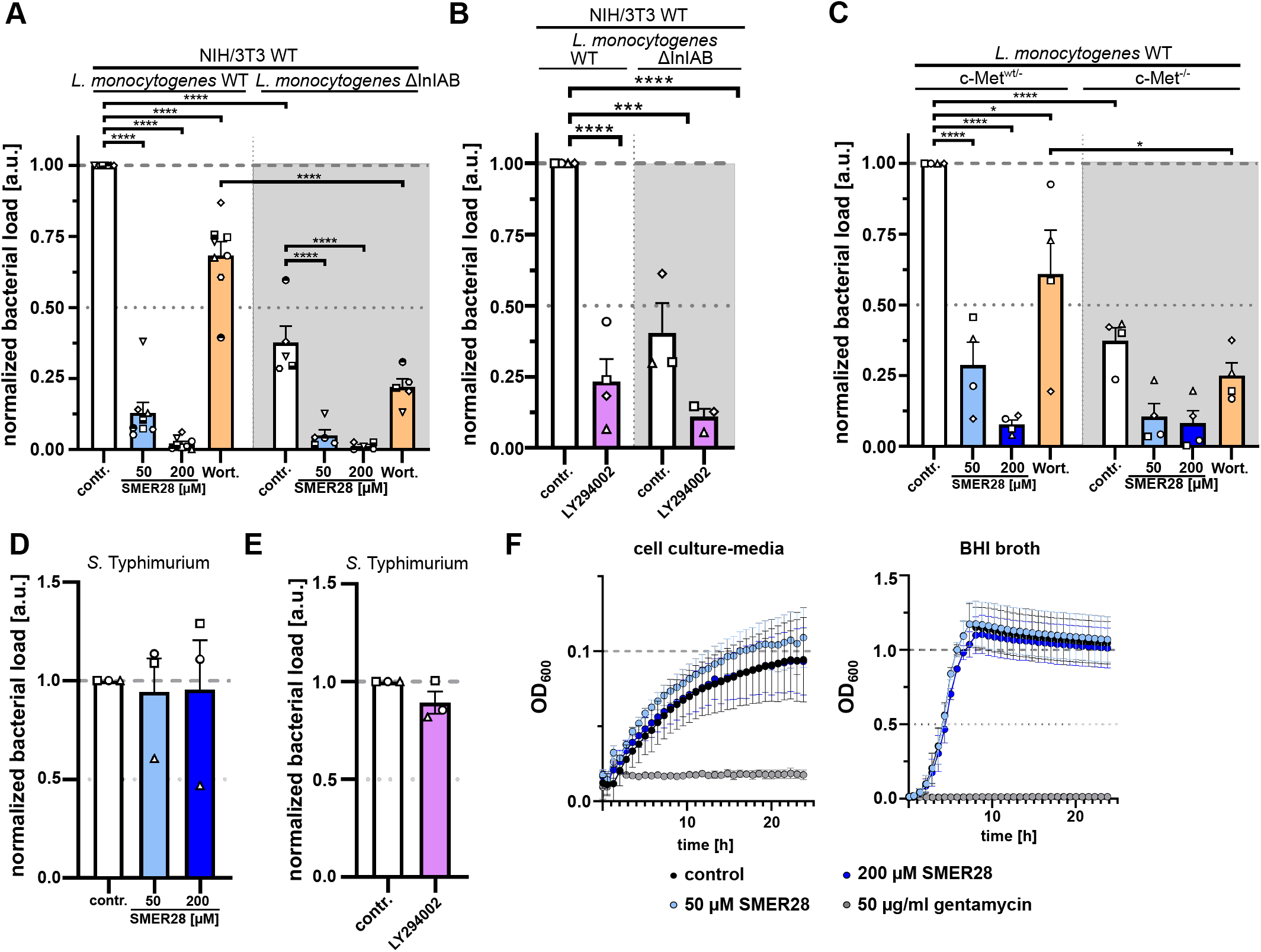
SMER28 efficiently inhibits *Listeria* uptake. **(A)** NIH/3T3 cells pre-treated with or without 50 or 200 µM SMER28 or 50 nM wortmannin for 16h were explored by gentamycin protection assay for invasion of *L. monocytogenes* wildtype and ΔInIA/B mutant. **(B)** NIH/3T3 cells were pre-treated with or without 50 µM LY294002 for 30 minutes and subjected to gentamycin protection assay using *L. monocytogenes* wildtype and ΔInIA/B. **(C)** c-Met^WT/-^and c-Met^-/-^cells were pre-treated as described in A, and examined by gentamycin protection assay with *L. monocytogenes* wildtype as above. **(D)** NIH/3T3 cells were pre-treated with or without 50 or 200 µM SMER28 for 16h. Cells were then subjected to gentamycin protection assay with *S*. Typhimurium as invasive pathogen. (E) NIH/3T3 cells were pre-treated with or without 50 µM LY294002 for 30 minutes and examined by gentamycin protection assay for invasion of *S*. Typhimurium. In (A-E), all bars display means from at least three independent experiments ± s.e.m. **(F)** Planctonic growth assay of *L. monocytogenes* wildtype bacteria in cell culture media (left graph) and in BHI broth (right graph). Media were supplemented with or without 50 µM SMER28, 200 µM SMER28 or 50 µg/ml gentamycin, as indicated. OD600 was measured every 15 min for 24h. Graphs display means from three independent experiments ± s.e.m. *p<0.05, ***p<0.001****p<0.0001; one way ANOVA.

## Discussion

The initial reports describing an mTOR-independent, autophagy-inducing potential of SMER28 left an important question unanswered: what is the direct target of SMER28 mediating this activity (19)? While we were able to confirm a modest, autophagy-inducing effect of SMER28 here, we discovered, to our surprise, that SMER28 acts on the RTK signaling axis by directly binding to and inhibiting PI3K. More specifically, SMER28 exhibits its strongest inhibitory activity on functional class I PI3K complexes comprising the catalytic subunit p110 delta. We therefore propose to re-name SMER28 to delsulisib (delta subunit inhibitor) to allow identification of this drug as PI3K inhibitor. It remains to be discovered whether SMER28 may have additional, biological targets. The enzymatic activity of PI3K is connected to essential cellular processes such as cell growth, cell cycle regulation and cell migration (50). In this study, we are able to connect several cell biological phenomena to SMER28-mediated *in vitro* inhibition of PI3K. We show that SMER28 has a cytostatic effect that is accompanied by low cytotoxicity, highlighting the role of PI3K as a growth factor/nutrient sensor, since inhibition of PI3K in malignant cells is more commonly cytostatic than cytotoxic (51). Moreover, we find that SMER28 has immediate effects on dynamic changes of the cytoskeleton such as dorsal ruffle formation or long-lasting effects like cell scattering, both of which require cytoskeletal rearrangements (32); such types of plasticity changes are key characteristics of cancer cells.

Aberrant PI3K signaling is commonly occurring in many types of cancers. Any type of misregulation of PI3K activity may lead to hyperactivation of the pathway, either by hyperactivating mutations of the enzyme itself, increased expression, increased PI3K copy numbers or loss of PI3K suppressors such as its direct antagonist PTEN. This often contributes to the development of diseases, such as cancer, but also immunological or neurological disorders as well as diabetes (5, 6, 18, 51). Thus, the development of therapeutics for treatments based on PI3K has been driven forward with much effort. These activities resulted in identification of a number of drugs that can be subdivided into dual PI3K/mTOR inhibitors, pan-PI3K inhibitors and isoform-specific inhibitors (16). Class I PI3K comprises four different members that are differentiated by their catalytic subunit p110. PI3Kα and PI3Kβ are expressed ubiquitously, while PI3Kγ and PI3Kδ are enriched in the hematopoietic system (5). B-cell malignancies such as multiple myeloma (MM), chronic lymphocytic leukemia (CLL) and indolent Non-Hodgkin Lymphoma (iNHL) are strikingly dependent on PI3Kδ activity (51, 52). Hence, drugs specifically targeting this enzyme were in demand for treatment of these lymphomas. The first PI3Kδ-specific inhibitor idelalisib (CAL-101) was approved by the FDA in 2014 (53-55). Notably, idelalisib induces autophagy (52). Unfortunately, idelalisib-treatment of relapsed CLL is accompanied by hepatotoxicity (56), which is hypothesized to be an off-target effect based on its structure. Subsequently-developed PI3Kδ-selective inhibitors such as parsaclisib and umbralisib were thus modified but based on the chemical structure of idelalisib (57). Although the chemical structure of SMER is known (19), the identification of the structural basis SMER28 binding to p110 delta will certainly help to understand the exact mechanism of inhibition. Moreover, it will be interesting to address whether the low cellular cytotoxicity of SMER28 observed here could be translated into human application. Strikingly, the systemic treatment of mice with SMER28 (58) fitted the low cytotoxicity that had been reported earlier (59, 60) and could be confirmed here, thus constituting promising insight for future, organismic application. Specifically, our findings imply that SMER28 might be an applicable drug for treatment of B-cell Non-Hodgkin-lymphoma. In addition to this, the PI3K signaling pathway emerges as central switch for effective infection of diverse pathogenic viruses such as ebola virus (61), West Nile virus (62) and Kaposi’s sarcoma herpesvirus (63). Notably, APDS (activated PI3Kδ syndrome), an inherited immune disorder, is characterized by susceptibility to herpesviruses, such as Epstein-Barr-Virus (EBV) (64). Importantly, inhibition of PI3Kδ is currently even discussed as potential target to prevent SARS-CoV-2 infection through suppression of the release of pro-inflammatory cytokines (65). Further exploration of this hypothesis promises to provide important, potential knowledge for defeating infectious diseases in the future. Finally, our study also illustrates how different signaling pathways are intertwined with each other and sheds light on the connections between RTK signaling and autophagy.

## Materials and Methods

### Antibodies and chemical compounds

Antibodies, chemical compounds and other materials are detailed in the supplementary reagents and materials table.

### Bacterial strain and cell line culture

*Salmonella enterica* serovar Typhimurium strain SL1344 was grown at 37°C in Luria-Bertani (LB) medium under agitation. *Listeria monocytogenes* (EGD [wild type]) and its isogenic internalin A and B-deleted strain (ΔInlAB) were grown at 37°C in brain heart infusion agar (BHI) medium under agitation.

U-2 OS human osteosarcoma cells, NIH/3T3 mouse embryonic fibroblast cells, c-Met control (WT/-) and KO (-/-) cells and Madin-Darby canine kidney (MDCK) cells were cultured in growth medium as follows: DMEM (4.5 g l^-1^ glucose) with 10% FCS, 2 mM L-glutamine, 1 mM sodium pyruvate and 1% non-essential amino acids. Cells were incubated at 37°C in a humidified 7.5% CO_2_-atmosphere. C-Met control (genotype WT/-) and KO fibroblast cells were individually harvested from embryos of breedings with mice heterozygous for a c-Met null allele (66). Upon genotyping, primary cultures with genotype WT/-as control and -/- were immortalized by retroviral transduction of SV40 LT antigen (67), using standard protocols.

### Cell lysate, Protein Measurements and Western Blotting

Protein extracts were prepared by harvesting cells that reached a confluency of approx. 70%. Cells were washed thrice with ice-cold PBS and lysed in ice-cold lysis buffer (50 mM Tris, pH 7.5, 150 mM NaCl, 1 mM EDTA, 1% Triton-X 100; Figure 1D, E and Figure supplement 1B) and SDS-laemmli sample buffer (Figure 3C-E, and 5A), respectively. The protein concentrations of cleared supernatants were detemined using a BCA Protein Assay kit. Proteins were separated by SDS-PAGE and blotted according to standard procedures. Primary antibodies used in this study are listed in the key resource table. Signals were detected by using the Lumi-Light Western Blotting Substrate (Roche) with the ECL Chemocam IMAGER (Intas, Göttingen, Germany). Densitometric analyses of respective proteins were normalized to the levels of GAPDH, Tubulin or total protein, as indicated.

### Propidium iodide cytotoxicity assay

U-2 OS cells were seeded into IncuCyte ImageLock 96-well plates (EssenBioscience) at a density of 3.000 cells/well, followed by replacement of the growth medium after 6 h with medium containing 0.5 µM propidium iodide and respective treatment as indicated. Cells were analyzed in an Incucyte S3 live-cell analysis system (EssenBioscience) with a 10x objective at 37°C and 5% CO_2_. Every 30 min, one image per well was collected by phase-contrast and red fluorescence for a time period of 47 h. Quantifications of cell confluence and PI-positive cell area were performed with the Basic Analyzer (Incucyte), and normalized to initial values.

### Cell proliferation assay

U-2 OS cells were seeded into a 24 well plate (3524, Corning) at a density of 20.000 cells/well and incubated for 16 hours. Media were replaced with medium containing DMSO alone as vehicle control or 50 µM or 200 µM SMER28 or 300 nM rapamycin. Cells were analyzed in an Incucyte S3 live-cell analysis system with a 10x objective at 37°C and 5% CO_2_. Four phase contrast images per well and four wells per treatment were acquired every hour for 47 hours. Quantification of cell confluence was carried out with the Basic Analyzer (Incucyte) and normalized to the average, respective initial confluence.

### Cell Cycle Analysis

For cell cycle analysis, U-2 OS cells were treated for 24 hours as indicated. Cells were harvested by centrifugation and pellets washed twice with ice cold PBS. Cells were fixed with 70% of ice-cold ethanol for 30 min followed by washing with PBS twice. 50 µg/ml RNase A and 50 µg/ml propidium iodide solution was added to remaining cell suspensions. Subsequently, cell fluorescence was measured using the BD LSRII SORP system (BD Bioscience). Fluorescence intensity analyses were performed with FlowJo software (BD Biosciences).

### Cell Scattering Assay

For scattering experiments, MDCK cells were seeded into a 24 well plate at a density of 3000 cells/well. Cells were stimulated after 32 h with 20 ng/ml HGF with or without 50 µM or 200 µM SMER28 as well as DMSO as vehicle control. Cell scattering was imaged every 15 minutes for 12h by phase contrast microscopy using a 10x objective at 37°C in a humidified 7.5% CO_2_-atmosphere. For analysis of MDCK colony scattering after 12h, scattered colonies were identified as cells that had lost contact to their neighboring cells.

### Growth factor response assay

For growth factor-induced membrane ruffling, 3×10^4^ NIH/3T3 cells were seeded onto fibronectin-coated glass coverslips in 24-well plates. Cells were washed with PBS three times and starved with DMEM containing respective inhibitors or vehicle control for 16 hours. Then, cells were stimulated with 20 ng/ml HGF or 10 ng/ml PDGF in DMEM for 5 minutes as indicated. Coverslips were washed twice with cytoskeleton buffer (CB) (10 mM MES, 150 mM NaCl, 5 mM EGTA, 5 mM glucose, 5 mM MgCl_2_, pH 6.1) and fixed with pre-warmed 4% paraformaldehyde (PFA) in cytoskeleton buffer. Cells were permeabilized with 0.1% Triton X-100 in CB for 1 min, and stained for the actin cytoskeleton with phalloidin Alexa Fluor 488 in CB for 1h at room temperature. All coverslips were washed in CB and mounted with ProLong Diamond Antifade.

### PI3 Kinase inhibitor assay

In vitro PI3 Kinase activity measurements were performed with the PI3K Kinase Activity/Inhibitor assay kit following manufacturer’s instructions (Merck Millipore). Briefly, kinase reactions were performed in provided glutathione-coated strips. Recombinant proteins of the isoforms of PI3K p110α, -β, -γ and -δ were each pre-incubated with DMSO control, 50 µM and 200 µM SMER28 for 10 min. Kinase reaction buffer, PIP_2_ substrate and distilled H_2_O were added, followed by incubation at room temperature for 1 h. Biotinylated-PIP_3_/EDTA solution was added to all wells except the buffer control. Then, GRP1 solution was added and incubated for 1 h at room temperature. After washing 4 times with TBST, SA-HRP solution was added and incubated for one more hour. Next, the wells were washed 3x with 1xTBST and twice with TBS before substrate TMB solution was added to each well. The reaction was incubated for 20 minutes in the dark and stopped by adding stop solution. Absorbance was read at 450 nm (Tecan Infinite 200pro).

### Gentamycin protection assay

For *Salmonella* Typhimurium and *Listeria monocytogenes* invasion assays, 5×10^4^ cells per well were seeded into a 24-well plate 24h before infection and incubated at 37°C in a humidified 7.5% CO_2_ atmosphere. Cells were pre-treated as indicated in figure legends. For *Salmonella* Typhimurium, LB broth was inoculated with an overnight culture (1:50) and grown at 37°C under agitation and an OD_600_ of 0.6-0.8. Bacteria were harvested by centrifuging at 3,000 x g for 3 min, and the bacterial density adjusted to a multiplicity of Infection (MOI) of 100 in DMEM. Infections of respective cell lines were initiated after replacing media with pre-warmed DMEM, adding Salmonella in DMEM and centrifugation for 5 min at 935 x g. After incubation for 30 min, 50 µg/ml gentamycin in DMEM was added for 30 min.

For *Listeria* gentamycin protection assays, the overnight culture was directly harvested by centrifuging at 3,000 x g for 3 min and the bacterial density adjusted to a MOI of 100 in DMEM. Infection of respective cell lines was performed essentially as described above, except that after incubation for 60 min, cells were washed once with PBS before addition of 50 µg/ml gentamycin in DMEM for 2 hours. Cells were then washed three times with PBS and lysed with 0.5% Triton X-100 in PBS for 5 min at room temperature. Lysates of all conditions were harvested on ice, and serial dilutions in PBS plated onto agar plates, followed by incubation at 37°C overnight. Colonies were then counted with an automatic colony counter (Scan4000, Interscience).

### Planktonic bacterial growth assay

Growth analysis of *L. monocytogenes* was performed with the Bioscreen C MBR. *L. monocytogenes* was grown at 37°C in BHI broth overnight under agitation. The OD_600_ was measured and set to an OD_600_ of 0.01 in BHI broth and fibroblast media supplemented with 50 µM SMER28, 200 µM SMER28 or 50 µg/mL gentamycin. OD_600_ was measured in quadruplicates every 15 minutes for 24 h during continuous shaking at 37°C using EZExperiments.

### Immunofluorescence

For immunofluorescence analysis, cells were seeded onto ethanol/acid-washed glass coverslips that were coated with 25 µg/ml fibronectin in PBS for 60 min at RT essentially as described previously (68). U-2 OS cells were cultured overnight before treatments were applied. Cells were fixed with iced methanol (−20°C) for 5 min and sequentially rehydrated in PBS. After PBS washing, the cells were blocked with 5% horse serum in PBS containing 0.3% Triton X-100 for 30 min. Upon blocking, cells were incubated in primary antibody solution in 1% BSA in PBS for 1 hour, washed with PBS and labeled with secondary antibodies for 40 min. After washing with PBS, cells were mounted with ProLong Diamond Antifade.

### Image acquisition

Fluorescence images shown in Figure 1A were acquired on a CSU-X1 (Yokogawa) spinning disk confocal microscope using a quad band filter connected to a Nikon Ti-Eclipse and a Modular Laser System 2.0 (Perkin Elmer) equipped with 405, 488, 561 and 640 laser lines. Plan Apochromat immersion objectives 60x oil/1.4 NA and 100x oil/1.4 NA (Nikon) and an EMCCD C9100-02 camera (Hamamatsu) were used. The system was controlled by the acquisition software Volocity 6.2.1 (Perkin Elmer). The fluorescence images shown in Figure 4C were acquired by a Nikon Ti2-Eclipse microscope with the spinning disk confocal module CSU-W1 (Yokogawa) using BP filters. Images were acquired with a Plan Fluor 40x oil/NA1.3 objective (Nikon), a Zyla 4.2 sCMOS camera (Andor) and 405/488/561/638nm laser lines (Omicron, Germany) controlled by NIS-Elements software. For 3D structured illumination microscopy (SIM), cells were imaged with a Nikon Ti-Eclipse Nikon N-SIM E microscope and a CFI Apochromat TIRF 100x Oil/NA 1.49 objective (Nikon). Image acquisition was controlled by the NIS-Elements software controlling an Orca flash 4.0 LT sCMOS camera (Hamamatsu), a Piezo z drive (Mad city labs), a LU-N3-SIM 488/561/640 laser unit (Nikon) and a motorized N-SIM quad band filter with a separate 525/50 emission filter using a 488 laser at 100% output power. Z-stack images were acquired with a step size of 200 nm. Reconstruction was carried out with the slice reconstruction tool (NIS-Elements, Nikon) using reconstruction parameters IMC 2.11, HNS 0.37, OBS 0.07.

### Data Processing and Statistical Analyses

Image analysis was carried out using ImageJ, Metamorph and NIS-Elements. Quantifications of LC3 and p62 structures were conducted by using the particle analyzer tool. Further data processing steps and statistical analyses were carried out in NIS-Elements, ImageJ, Inkscape, Excel 2010, and Graphpad Prism 9. Results from statistical tests, sample sizes and number of experiments are given in respective figure legends.

## Supporting information

Reagents and materials table

## Acknowledgements

We would like to thank Peer Lukat (HZI) for fruitful discussions. This research was supported in part by the Deutsche Forschungsgemeinschaft (to KR, individual grant RO 2414/8-1) and by the Helmholtz Society through intramural funding as well as the HGF impulse fund W2/W3-066TEBS (to TEBS).

## Figure Legends

**Figure 1 - Supplement Figure 1**

**SMER28 stimulates autophagy in U-2 OS cells. (A)** Quantifications of the areas of LC3- and p62-positive puncta per autophagosome. Scatter plots from three independent experiments. Data represent means ± s.e.m.; n= total number of cells analyzed. **(B)** U-2 OS cells were treated with or without 50 µM SMER28, 1 µM MG-132, 15 nM bortezomib (BZE) and in the presence or absence of 250 nM bafilomycin A1 or 300 nM rapamycin for 16 h, as indicated. Samples were probed for poly-ubiquitinated proteins.

**Supplementary Movie 1. (A)** HGF-induced cell scattering of MDCK cells with pre-formed colonies. Cells were serum-starved in DMEM with or without 200 µM SMER28 for 16h, as indicated. Cells were stimulated with 20 ng/ml HGF in respective treatment and imaged every 30 min for 12h. Scale bar: 100 µm.

