## Supplementary figures and images for "SMER28 attenuates PI3K/mTOR signaling by direct inhibition of PI3K p110 delta"

### Supplement Figure 1

#### Figure 1 - supplement

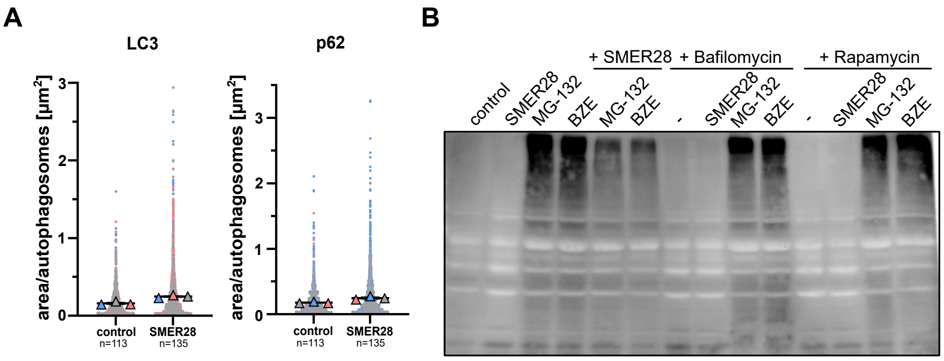
