## Supplementary material for "SMER28 attenuates PI3K/mTOR signaling by direct inhibition of PI3K p110 delta": Reagents and materials table

| Reagent type (species) or resource | Designation | Source or reference | Identifiers | Additional Information |
|---|---|---|---|---|
| Antibody | Anti-LC3A/B (Rabbit polyclonal) | Cell Signaling Technology, Beverly, MA | 4108S RRID: AB_2137703 | 1:5000 |
| Antibody | Anti-mTOR (7C10) (Rabbit monoclonal) | Cell Signaling Technology, Beverly, MA | 2983 RRID: AB_2105622 | 1:2000 |
| Antibody | Anti-phospho-mTOR (Ser2448) (D9C2) (Rabbit monoclonal) | Cell Signaling Technology, Beverly, MA | 5536 RRID: AB_10691552 | 1:2000 |
| Antibody | Anti-Akt (Rabbit polyclonal) | Cell Signaling Technology, Beverly, MA | 9272 RRID: AB_329827 | 1:2000 |
| Antibody | Anti-phospho-Akt (Ser473) (Rabbit polyclonal) | Cell Signaling Technology, Beverly, MA | 9271 RRID: AB_329825 | 1:2000 |
| Antibody | Anti-phospho-Akt (Thr308) (Rabbit polyclonal) | Cell Signaling Technology, Beverly, MA | 9275 RRID: AB_329828 | 1:2000 |
| Antibody | Anti-phospho-PI3 Kinase p85 alpha (Tyr607) (Rabbit polyclonal) | Abcam, Cambridge | ab182651 RRID: AB_2756407 | 1:2000 |
| Antibody | Anti-PI3-Kinase (Mouse monoclonal) | BD Transduction Laboratories | 610045 RRID: AB_397459 | 1:2000 |
| Antibody | Anti-phopho-ULK1 (Ser757)(D7O6U) (Rabbit monoclonal) | Cell Signaling Technology, Beverly, MA | 14202 RRID: AB_2665508 | 1:2000 |
| Antibody | Anti-ULK1 (D8H5) (Rabbit monoclonal) | Cell Signaling Technology, Beverly, MA | 8054 RRID: AB_11178668 | 1:2000 |
| Antibody | Anti-phospho-4E-BP1 (Thr37/46) (236B4) (Rabbit monoclonal) | Cell Signaling Technology, Beverly, MA | 2855 RRID: AB_560835 | 1:2000 |
| Antibody | Anti-4E-BP1 (Rabbit polyclonal) | Cell Signaling Technology, Beverly, MA | 9452 RRID: AB_331692 | 1:2000 |
| Antibody | Anti-phospho-p70 S6 Kinase (Thr389) (Rabbit polyclonal) | Cell Signaling Technology, Beverly, MA | 9205 RRID: AB_330944 | 1:2000 |
| Antibody | Anti-p70 S6 Kinase (Rabbit polyclonal) | Cell Signaling Technology, Beverly, MA | 9202 RRID: AB_331676 | 1:2000 |
| Antibody | Anti-SQSTM1/p62 (Mouse monoclonal) | Abcam, Cambridge | ab56416 RRID: AB_945626 | 1:2000 |
| Antibody | Anti-Ubiquitin (P4D1) (Mouse monoclonal) | Cell Signaling Technology, Beverly, MA | 3936 RRID: AB_331292 | 1:2000 |
| Antibody | Anti-GAPDH (6C5) (Mouse monoclonal) | Calbiochem | CB1001 RRID: AB_2107426 | 1:5000 |
| Antibody | Anti-Tyr-Tubulin (Clone YL 1/2) (Rat monoclonal) | Jürgen Wehland | | 1:2000 |
| Antibody | Anti-p21 Waf1/Cip1 (12D1) (Rabbit monoclonal) | Cell Signaling Technology, Beverly, MA | 2947 RRID: AB_823586 | 1:2000 |
| Antibody | Anti-p27 Kip1 (D69C12) XP (Rabbit monoclonal) | Cell Signaling Technology, Beverly, MA | 3686 RRID: AB_2077850 | 1:2000 |
| Antibody | Anti-Cyclin D1 (92G2) (Rabbit monoclonal) | Cell Signaling Technology, Beverly, MA | 2978 RRID: AB_2259616 | 1:2000 |
| Antibody | Anti-Cyclin D3 (DCS22) (Mouse monoclonal) | Cell Signaling Technology, Beverly, MA | 2936 RRID: AB_2070801 | 1:2000 |
| Antibody | Anti-CDK4 (D9G3E) (Rabbit monoclonal) | Cell Signaling Technology, Beverly, MA | 12790 RRID: AB_2631166 | 1:2000 |
| Antibody | Anti-CDK6 (DCS83)(Mouse monoclonal) | Cell Signaling Technology, Beverly, MA | 3136 RRID: AB_2229289 | 1:2000 |
| Antibody | Anti-mouse IgG (H+L)-HRPO (Goat polyclonal) | Jackson ImmunoResearch, Cambridge | 115-035-062 RRID: AB_2338504 | 0.16 mg/ml |
| Antibody | Anti-rabbit IgG (H+L)-HRPO (Goat polyclonal) | Jackson ImmunoResearch, Cambridge | 111-035-045 RRID: AB_2337938 | 0.16 mg/ml |
| Antibody | Anti-rat IgG (H+L)-HRPO (Goat polyclonal) | Jackson ImmunoResearch, Cambridge | 112-035-062 RRID: AB_2338133 | 0.16 mg/ml |
| Antibody | Anti-mouse IgG (H+L)-Alexa Fluor 594 (Goat polyclonal) | Invitrogen (Molecular Probes) | A11032 RRID: AB_2534091 | 1:400 |
| Antibody | Anti-rabbit IgG (H+L)-Alexa Fluor 488 (Goat polyclonal) | Invitrogen (Molecular Probes) | A11034 RRID: AB_2576217 | 1:400 |
| Antibody | Anti-LC3B (Rabbit polyclonal) | Novus Biologicals | NB600-1384 RRID: AB_669581 | 1:500 |
| Chemical compound, drug | Phalloidin-Alexa Fluor-488 | Thermo Fisher Scientific | A12379 | 2.25 U/ml |
| Chemical compound, drug | Propidium Iodide | BioLegend | 421301 | |
| Chemical compound, drug | Bortezomib | Cayman Chemicals, Ann Arbor, MI | CAS 179324-69-7 | dissolved in DMSO |
| Chemical compound, drug | MG-132 | Cayman Chemicals, Ann Arbor, MI | CAS 133407-82-6 | dissolved in DMSO |
| Chemical compound, drug | Rapamycin | Cayman Chemicals, Ann Arbor, MI | CAS 53123-88-9 | dissolved in DMSO |
| Chemical compound, drug | Bafilomycin A1 | Cayman Chemicals, Ann Arbor, MI | CAS 88899-55-2 | dissolved in DMSO |
| Chemical compound, drug | Epothilone B | Mark Stadler, HZI, Braunschweig | (040_006a) | dissolved in DMSO |
| Chemical compound, drug | Paclitaxel | StressMarq Biosciences, Victoria, Canada | SIH-239 | dissolved in DMSO |
| Chemical compound, drug | Wortmannin | ALEXIS Biochemicals | 350-202-M001 | dissolved in DMSO |
| Chemical compound, drug | Small molecule enhancer of rapamycin 28 (SMER28) | Sigma-Aldrich | S8197 (Lot #043M4613V) | dissolved in DMSO |
| Chemical compound, drug | LY294002 | Sigma-Aldrich | 19-142 | dissolved in DMSO |
| Chemical compound, drug | Gentamycin | Sigma-Aldrich | G1272 | |
| Commercial assay or kit | PI3 Kinase Activity/Inhibitior ELISA | Sigma-Aldrich | 17-493 | |
| Commercial assay or kit | BCA Protein Assay Kit | Pierce | 23225 | |
| Other | Fetal Bovine Serum (FBS) | Sigma-Aldrich | F7524 (Lot #05443396) | |
| Other | Dulbecco's Modified Eagle Medium (DMEM) | Gibco | 41965-039 | |
| Other | ProLong™ Diamond Antifade Mountant | Invitrogen | P36962 | |
| Other | PBS | Gibco | 10010015 | |
| Other | HGF | Sigma-Aldrich | H9661 | |
| Other | RNase A | Thermo Fisher Scientific | EN0531 | |
| Other | HBSS | Biochrom | L2035 | |
| Other | PDGF | Sigma-Aldrich | P3201 | |
| Other | Precision cover glasses 12 mm thickness No. 1.5H | Paul Marienfeld | 117580 | |
| Other | Glass slide | Thermo Fisher Scientific | AAAA000082##32E | |
| Other | Fibronectin | Roche | 11051407001 | 25 µg/ml in PBS |
| Other | IncuCyte ImageLock 96-well plates | EssenBioscience | 4379 | |
| Other | Trition X-100 | Bio-Rad | 1610407 | |
| Other | Honeycomb 2 plates | Bioscreen | 95025BIO | |
| Other | Bacto Brain Heart Infusion | Becton Dickinson | 237500 | |
| Other | Bacto Agar | Becton Dickinson | 214010 | |
| Other | Bacto Yeast Extract | Becton Dickinson | 212750 | |
| Cell line (*Homo sapiens)* | U-2 OS | DSMZ | ACC785 | |
| Cell line (*M. musculus)* | NIH/3T3 | ATCC | CRL-1658 | |
| Cell line (*Canis familiaris)* | MDCK | ATCC | CCL-34 | |
| Bacteria | Salmonella enterica serovar Typhimurium wildtype SL1344 | Hoiseth & Stocker, Nature, 1981 | | doi: 10.1038/291238a0. |
| Bacteria | Listeria monocytogenes EGD [wild type] | Kaufmann, Infection and Immunity, 1984 | | doi: 10.1128/iai.45.1.234-241.1984 |
| Bacteria | Listeria monocytogenes) ∆InIAB | Parida et al., Mocelular Microbiology, 2002 | | doi.org/10.1046/j.1365-2958.1998.00776.x |
| Software, algorithm | Image J | NIH; Schneider et al., 2012 | https://imagej.nih.gov/ij/ | |
| Software, algorithm | Prism 9 | Graphpad | https://www.graphpad.com/ | |
| Software, algorithm | MetaMorph | Molecular Devices | https://www.metamorph.com/ | |
| Software, algorithm | NIS-Elements | Nikon Instruments | https://www.microscope.healthcare.nikon.com/en_EU/products/software/nis-elements | |
| Software, algorithm | FlowJo™ | BD Biosciences | https://www.flowjo.com/ | |
| Software, algorithm | Inkscape | Inkscape Project | https://inkscape.org | |
